# Harmonization techniques for machine learning studies using multi-site functional MRI data

**DOI:** 10.1101/2023.06.14.544758

**Authors:** Ahmed El-Gazzar, Rajat Mani Thomas, Guido van Wingen

## Abstract

In recent years, the collection and sharing of resting-state functional magnetic resonance imaging (fMRI) datasets across multiple centers have enabled studying psychiatric disorders at scale, and prompted the application of statistically powerful tools such as deep neural networks. Yet, multi-center datasets introduce non-biological heterogeneity that can confound the biological signal of interest and produce erroneous findings. To mitigate this problem, the neuroimaging community has adopted harmonization techniques previously proposed in other domains to remove site-effects from fMRI data. The reported success of these approaches in improving the generalization of the models have varied significantly. It remains unclear whether harmonization techniques could boost the final outcome of multi-site fMRI studies, to what extent, and which approaches are best suited for this task. In an attempt to objectively answer these questions, we conduct a standardized rigorous evaluation of seven different harmonization techniques from the neuroimaging and computer vision literature on two large-scale multi-site datasets (*N* = 2169 and *N* = 2366) to diagnose autism spectrum disorder and major depression disorder from static and dynamic representations of fMRI data. Interestingly, while all harmonization techniques removed site-effects from the data, they had little influence on disorder classification performance in standard k-fold and leave-one-site-out validation settings over a well-tuned baseline. Further investigation shows that the baseline model implicitly learns site-invariant features which could well explain its competitiveness with explicit harmonization techniques and suggest orthogonality between latent disease features and site discrminative features. However, additional experiments show that harmonization methods could be critical to report faithful results in settings where there is high intra-site class imbalance and the learning algorithm is prone to overfit on spurious features confounding the final outcome of the study.

## 1. Introduction

Clinical machine learning (ML) studies on fMRI data, where models are trained to predict the diagnosis of brain disorders from resting-state functional activity, are at the core of recent research efforts in the field of psychiatry in an attempt to understand the pathophysiology of mental diseases, find objective biomarkers and improve the robustness of diagnosis in psychiatry. A central hurdle in this direction is the limited sample size a research group can collect at a single site to study a certain disorder. This hinders the generalization of the results and prevents the application of more statically powerful, yet less sample-efficient learning methods such as deep neural networks. In order to mitigate this problem, many researcher groups collaborate to create larger datasets, formed by aggregating fMRI scans obtained at different locations into a single multi-site dataset targeting a specific psychiatric disorder [1, 2, 3]. Unfortunately, inter-scanner variability, possibly caused by field strength of the magnet, manufacturer and parameters of the MRI scanner or radio-frequency noise environments [4, 5], creates a second problem known as batch effects, which is technical noise that might confound the real biological signal. The main consequence of batch effects is that researchers have observed a decrease in performance of the ML models on the larger multi-site datasets compared with performance obtained when using a dataset collected at a single site [6, 7]. Various approaches have been recently proposed to remove site-effects in the input fMRI signal post-acquisition. For example, [8] applied the Combat [9] harmonization technique, originally proposed to remove batch effects in genetic data, to estimate and regress site effects in functional connectivity while preserving the biological signal. [10] investigated the application of multiple linear transformations on the functional connectivity matrices to harmonize the data. [11] adopted a travelling subject dataset to estimate and remove measurement bias and improve the signal to noise ratio of the data. While these methods reported improvements in the generalization performance for their respective datasets and learning algorithms, further adoption of these techniques by multiple multi-site ML studies have not shown the same success. Data harmonization therefore remains a central area of controversy in the neuroimaging community.

A major limitation of popular harmonization techniques used for fMRI data such as Combat and Covbat is that they are only constrained to static functional connectivity representations and can not be further applied to other fMRI imaging derivatives (e.g. 4D volumes, timecourses, spatio-temporal graphs). Such representations capture the dynamic nature of the signal which have been shown as a more suitable candidate for finding disease biomarkers versus pre-defined static correlations [12, 13]. Moreover, fMRI harmonization studies have only employed linear models or simple kernel-based methods as the baseline classifiers to evaluate the effectiveness of the proposed approaches. A sub-optimal choice of the learning model could influence the results, as the baseline model could be more prone to learn spurious correlations in the data, which might not be the case if a more powerful baseline is used in the first place. Finally, state-of-the-art harmonization methods [14, 15, 16] in other domains, have shifted to harmonizing data in the latent space while optimizing for the label of interest, thus harmonizing the discrimnative features in an end-to-end manner. This is in contrast to harmonizing the data in the input space as a pre-processing step as is the current practice with fMRI data. The application of such techniques have transferred to different medical domains including structural MRI[17], yet, their success in the fMRI domain is yet to be assessed. This is in general a common theme in the literature where the large majority of harmonization studies focus on structural MRI volumes and derivatives [18, 19, 17, 20]. The problem is different for fMRI data and we argue is more difficult to tackle. The dynamic and stochastic nature of fMRI, the high dimensionality of the data, lack of standardized learning architectures, high noise in the labels, and limited sample sizes deems the evaluation of different harmonization methods a very challenging task. However we conjecture that it is an essential step moving forward if we are to find generalizable biomarkers for psychiatric disorders.

In this work, our aim is to build a reproducible standardized test-bed to measure and compare the effect of applying different harmonization techniques on static and dynamic representations of the fMRI data while using powerful deep neural networks for end-to-end diagnosis of psychiatric disorders. Namely, we implement 7 different harmonization techniques from the computer vision and neuroimaging literature (including ComBat, CovBat, Domain adversarial methods and deep feature harmonization) across two multi-site datasets on both static and dynamic fMRI representations using suitable choices of deep neural network architectures. The motivation to use deep learning models in our frame-work is i) the flexibility of neural networks (NN) to approximate any function and to incorporate different objectives simultaneously, enabling the implementation of several modern harmonization techniques. ii) The powerful representations capacity of NN as baseline models to enable a fair evaluation of the advantage of harmonization methods. iii) A main motivation behind data-pooling is the ability to use powerful tools like deep neural networks. Given the rapid developments of research efforts in building deep learning models for diagnosing psychiatric disorders[21], we believe it would be more useful to the community, especially since we designed our framework in an architecture-agnostic manner, and one could easily swap the neural networks used in the framework with a customized architecture.

In addition to the challenging nature of the data, we acknowledge that it is difficult to systematically compare different harmonization techniques given the stochastic nature of training neural networks and the variability in models sizes introduced by additional modules for some harmonization techniques. We strive for a fair comparison by standardizing the model architecture across methods, conducting extensive hyperparameter search for each method, and using multiple different seeds in running the experiments to report the best average test metrics under the best hyper-parameters for each method optimized on an independent validation set. This work was developed on top of DomainBed[22] framework to ensure a fair, reproducible and streamlined comparison of the algorithms against each other and against a strong well-tuned baseline. Our implementation is open-source and can be easily extended to incorporate and evaluate new harmonization methods, different phenotypes and different representations of fMRI data along with their respective learning architectures of choice.

We summarize our contributions as follows:

- First, we investigate the ability of the deep neural networks to classify sites from static and dynamic representations of the data. We conduct further experiments to understand how these effects are manifested in the imaging derivatives.
- We implement different harmonization algorithms from the neuroimaging and computer vision literature and conduct a rigorous evaluation to benchmark the utility of these algorithms in improving the generalizability of the trained model measured as improvement of phenotype prediction on leave-one-site-out and site-stratified k-fold validation schemes.
- We propose a simple empirical method to estimate site differences in the latent features used for classifcation of the biological label of interest.
- We study the effect of ML method selection, number of available training sites, and intra-site class imbalance on the generalization performance of harmonization methods.

## 2. Methods

### 2.1. Datasets and models

For the purpose of our analysis, we chose two publicly available multi-site datasets. Each one of the datasets targets a different psychiatric disorder, with a different demographic background and different pre-processing pipeline. This experimental setup eliminates results biases towards these variables and support the generalization of the empirical results across pre-processing pipelines, demographics, and learning objectives.

#### ABIDE I+II

contains a collection of rs-fMRI brain images aggregated across 29 institutions[1]. It includes 1,028 participants with a diagnosis of autism, Asperger or pervasive developmental disorder (called ASD from now on), and 1,141 typically developing participants (TD). In this study, we select the largest 9 sites in terms of number of participants (*N >* 50) to conduct our analysis. Figure 1 shows the distribution of participants between sites. Demographic information are reported in Table S1. The 4D volumes were pre-processed according to the CPAC pipeline [23](See supplementary materials for details of the pre-processing steps). Further, A craddock-200[24] atlas was used to parcellate the volume into 200 regions of interest (ROIs) and the timecourse of each region was calculated as the mean timecourse of all the voxels within the ROI.

**Figure 1:**
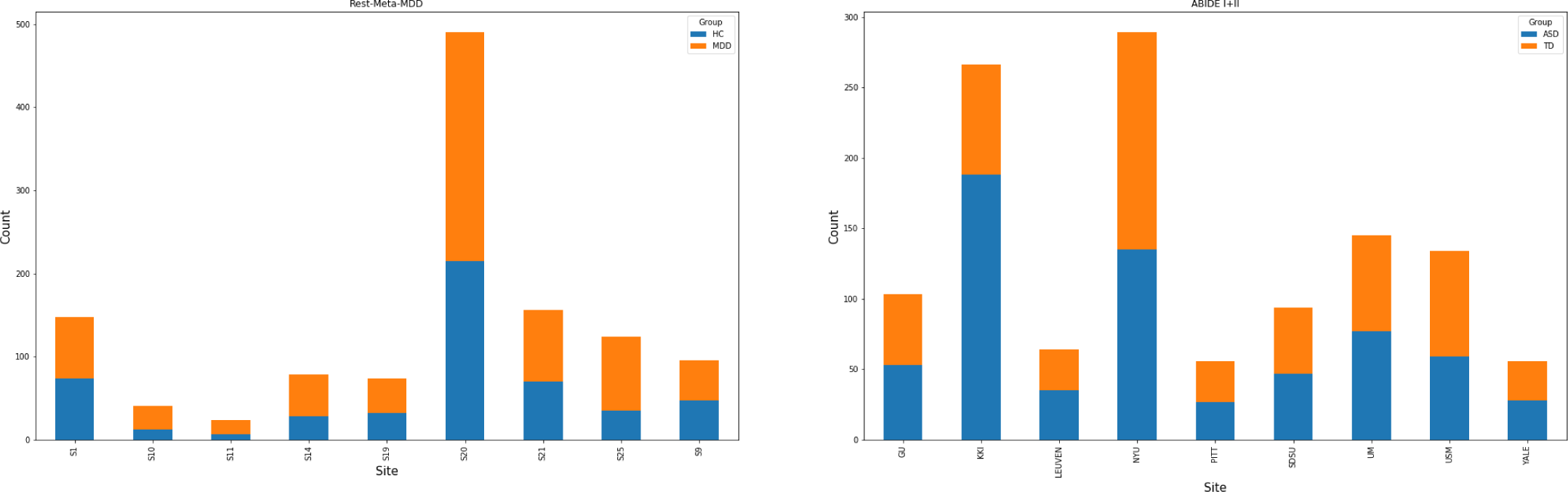
Site distribution of Rest-Meta-MDD and ABIDE I+II datasets.

#### Rest-Meta-MDD

is currently the largest resting-state fMRI database for studying major depression disorder (MDD) [25], including 1,255 patients and 1,083 HC from 25 cohorts in China. In this study we also select the largest 9 sites that contain at least 20 participants from each group. Figure 1B shows the distribution of participants between sites of the datasets. Demographic data are reported in Table S2 (see Supplementary Information for more details about sample composition). Standard preprocessing of the data was done at each site using the Data Processing Assistant for Resting-State fMRI (DPARSF) [26]. 116 time-courses of cortical and sub-cortical regions as defined by the Harvard-Oxford atlas [27] were extracted to obtain the input representations for the ML models.

### 2.2. Input representations and learning models

Another variable of interest in this study is the data representation used in the analysis. Different imaging derivatives can exhibit different levels of heterogeneity [28] and subsequently, display different results when evaluating the efficacy of harmonization methods. Thus, we chose two input representations to conduct our analysis, and for each representation we used a suitable learning model architecture.

#### Static - Pearson’s correlations

are the pairwise correlations between ROI timecourses. This results in a matrix of *N × N* matrix where *N* is the number of ROIs. The lower triangular matrix is then flattened and fed to the learning algorithm. This representation constitutes the most popular approach in ML in neuroimaging due to its simplicity, reduced dimensionality and the improvement of signal to noise ratio. In developing learning models for this representation, we chose multi-layer perceptrons (MLPs) []. Correlation vectors do not include any geometric inductive biases and thus fully connected layers offer a general statistical tool for feature extraction and classification, and have been successfully used for phenotype classification of correlations vectors [29]. Further, MLPs of-fer a flexible building block to implement different modern deep harmonization techniques as we further discuss in section 4.

#### Dynamic - ROI Time-courses

Unlike static correlations, feeding the input as the time-courses preserves the dynamic nature of the temporal signal. This can be challenging to model due to the sequential nature of the data and its high dimensionality coupled with the limited sample size of the dataset. A promising solution to model this signal is 1D convolutional neural networks (1D-CNNs)[30]. 1D-CNNs offer a computationally efficient approach to extract spatio-temporal features from fMRI data and have shown success in characterizing psychiatric disorders and phenotypes [31]. 1D-CNNs consists of a cascade of 1D convolutional blocks (1D Conv Layer[32] + Batch-Norm[33] + Max-Pooling + activation function) used for feature extraction followed by global average pooling and a fully connected layer for classifcation. Similarly, this model offers a flexible solution for architecture-agnostic implementations of deep harmonization techniques. Figure 2 offers a schematic of our learning pipelines.

**Figure 2:**
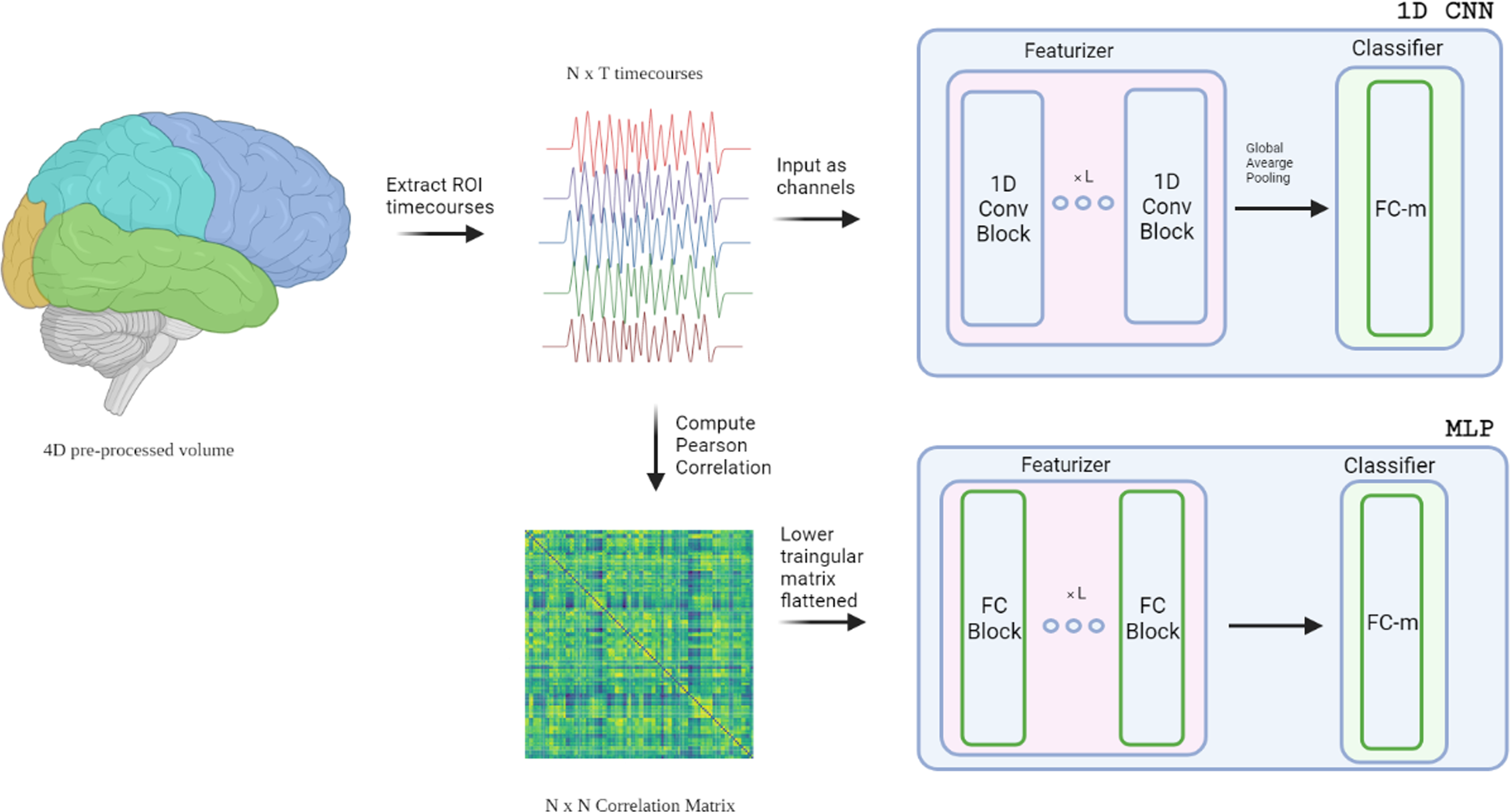
Learning pipelines for both static and dynamic representations of the data.

### 2.3. Site classification

Before delving into any harmonization methods, we first evaluated the performance of our pipelines in estimating the acquisition site on unseen data. The purpose of this experiment is three folds. 1) These results provide an empirical metric of heterogeneity in each dataset and each data representation. This metric can later be used to evaluate the efficacy of harmonization methods when applied and can provide a correlation measure of the effect of site heterogeneity on the performance of the main task in question. 2) Validate the ability of the proposed architectures to extract site-dependent features when supervised with the sites label as this is a perquisite for the implementation of some harmonization techniques. 3) Gain a better understanding of the nature of site differences in the imaging derivatives. (e.g. global vs local and linear vs non-linear)

To conduct this experiment, we trained the models as multi-class classifiers on the largest 4 sites in terms of number of samples in both datasets using both representations and their corresponding model architecture. Training details for both models is provided in the supplementary material. We report 5-fold cross-validation results on Table 1. The results show that acquisition site can be estimated with high accuracy (chance level = 25%) in both datasets. Further, it shows higher accuracy metrics when conducting the experiment on the timecourses indicating a higher heterogeneity in the dynamic signal over static connectivity. This makes intuitive sense as the temporal resolution of the timecourses is different across sites. This variability is absent in the static representations as the features only constitute the correlations of ROI signals. Another area of interest is the correlation of test metrics to the model’s linearity/non-linearity. This is evaluated using exclusively linear or non-linear activation functions after each feature extraction layer in the models. i.e. a linear 1D CNN only applies an identity transformation after each convolution layer while a non-linear 1D CNN applies a non-linear transformation (rectified linear unit here) after each convolution layer. Significantly higher test metrics when training using non-linear models would suggest that site differences are intrinsically non-linear and non-linear functions are more optimal to represent this relationship. However, this behaviour is not observed in the results as linear and non-linear models perform on-par on both datasets and using both representations. This could suggest that site differences manifest as a linear transformation and a linear classification model is sufficient to capture such differences. To further investigate if the reason for this high heterogeneity is local (i.e. a subset of features) or global, we conduct a simple ablation study, where only a subset of *N* regions is used to train and test the model. For each *N*, we randomly sample the regions included in the subset, conduct the site classification experiment, repeat the sampling and training 200 times, and report the mean test accuracy and standard deviations for each *N*. Results of this experiment conducted on the dynamic representations of the top-4 sites of the ABIDE dataset show that site effects can be captured (Acc. *≥* 30%) across any *N ≥* 5 regions in the signal, indicating the site effects is global across the input. Saliency maps of the trained MLP on the static representation also show high gradients across the entire signal indicating that site effects are global (See supplementary materials for figure and exact details to generate the saliency maps).

**Table 1:**
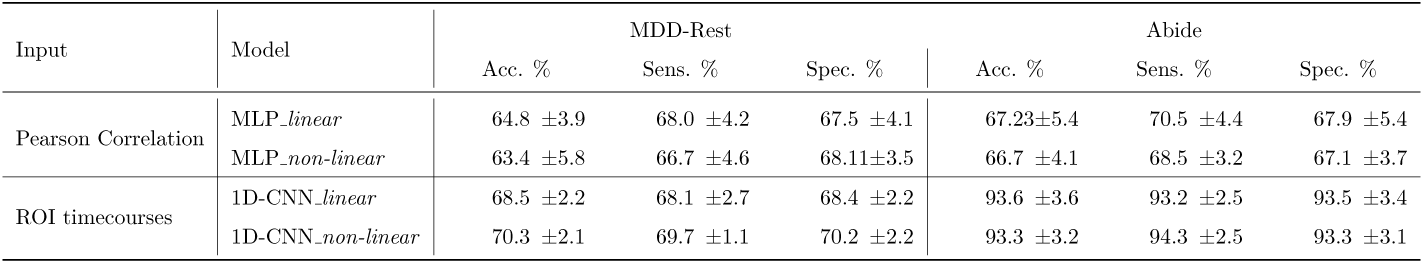
5-fold test results (mean *±* standard deviation) for site classification for the largest 4 sites in each dataset. Chance level is 25%.

### 2.4. Harmonization techniques

Strong site differences on the pre-processed imaging derivatives further confirm the existence of non-biological variability across sites and shows that standardized acquisition protocols and standardized pre-processing pipelines fail to eliminate such heterogeneity. This drives the need to find harmonization solutions to avoid erroneous findings. In this section we briefly describe some of the popular approaches in the computer vision and neuroimaging literature implemented in our framework. For further details about the methods, we refer the reader to the supplementary materials and the respective references of each method. We group the methods under two distinct categories and a baseline.

#### 2.4.1. Baseline: no explicit harmonization

To validate the efficacy of harmonization techniques, we conduct experiments where the model is trained on the pre-processed imaging derivatives without any explicit harmonization either in the input space or feature space and solely with the objective of minimizing the prediction error of the label of interest. A key aspect to ensure a fair evaluation of the baseline is to deploy equivalent computational resources spent on tuning the harmonization methods to find the most optimal baseline under the same computational time and budget. This for example include model size, training time, hyperparameter search space, data standardization, etc. Failing to guarantee a fair playground is a common pitfall in harmonization studies and can lead to overoptimistic conclusions about the efficacy of some harmonization methods in improving the final outcome over an under-powered baseline.

#### 2.4.2. Input harmonization

This group of techniques rely on the assumption that the distributional shift between sites is a linear transformation and could be mitigated via linear transformations of *X* into a shared harmonized input space. Common approaches in neuroimaging include z-scoring and whitening to remove batch effects caused by translation, scaling and rotation of input features. More general approaches to mitigate linear transformations that have been introduced and are quite popular in multiple medical and biological domains are ComBat[9] and CovBat[34]. Combat uses a multivariate linear mixed effects regression with terms for biological variables and scanner to model imaging feature measurements. The method uses empirical Bayes to simultaneously model and estimate biological and non biological terms and algebraically removes the estimated additive and multiplicative site effects. CovBat operates similarly to ComBat where in addition to correction of mean and variance across sites, it also corrects for covariance. Another recent approach from the computer vision and deep learning literature is Inter-domain Mixup[35], which performs linear interpolations between random pairs samples from different domains with the same label of interest. These interpolated examples are then fed to the learning model and the models are trained with no further explicit harmonization in the objective function. The intuition behind this approach is that the training distribution becomes siteagnostic while preserving the biological signal of interest presumably existing within data samples with the same label. In our analysis we include ComBat, CovBat and Inter-domain Mixup for the static input representation, and only Inter-domain Mixup to the the dynamic representation. Note that combat and covbat approaches can not be applied to dynamic timecourses given that there are no 1-to-1 associations between the input features across individual samples.

#### 2.4.3. Deep features harmonization

Recent advances in the domain generalization and domain adaption literature in deep learning rely on aligning the distribution of multiple domains in the latent space while training the network for the task of interest in an end-to-end manner. This also acts as a regulization technique and has been shown successful in multiple tasks. In this work we selected four of the most popular approaches for deep features harmonization.

- DANN [15]: Domain adversarial neural networks employ an adversarial network to match feature distributions across sites.
- IRM [16]: Independent Risk Minimization learns a feature representation *ϕ*(*X*) such that the optimal linear classifier on top of that representation matches across domains.
- CORAL [36]: matches the mean and covariance of feature distributions.
- MMD [14]: matches the maximum mean discrepancy of feature distributions.

## 3. Experiments

In order to evaluate the performance of a harmonization method, we would like answer two question in a quantitative manner; 1) *How successful is each harmonization method in removing site-induced differences?* 2)*What is the impact of applying a specific harmonization technique on the final outcome of the DL model?*

To answer the first question, we can revert back to the acquisition site prediction experiment we conducted on the imaging derivatives in Section 2.3 but now after harmonizing the data. The quantitative measure can thus be the difference between the test metric of the site classification preand postapplying harmonization. To recap, this means that we again *train* a new site classifcation model from scratch on the harmonized data and validate the performance using a 5-fold validation scheme. While this is straightforward for harmonization methods in the input space (e.g. ComBat, CovBat), it is more tricky for feature harmonization methods which harmonize the data in the feature space while training for the label of interest simultaneously. To enable conducting this experiment for these methods, we first train the models on the desired label of interest (while incorporating the harmonization method in the objective function), then only use the output of the trained featurizer to train the site classifier model without back-propagating the learning gradients of the sites through the featurizer. This means that the site classifier is trained only on the latent harmonized features and not the entire input data. We use the same approach to evaluate the features of the baseline method where we first train for the label of interest (without any explicit harmonization). Figure 3 illustrates the training process for the site classification experiment for both harmonized inputs and features.

**Figure 3:**
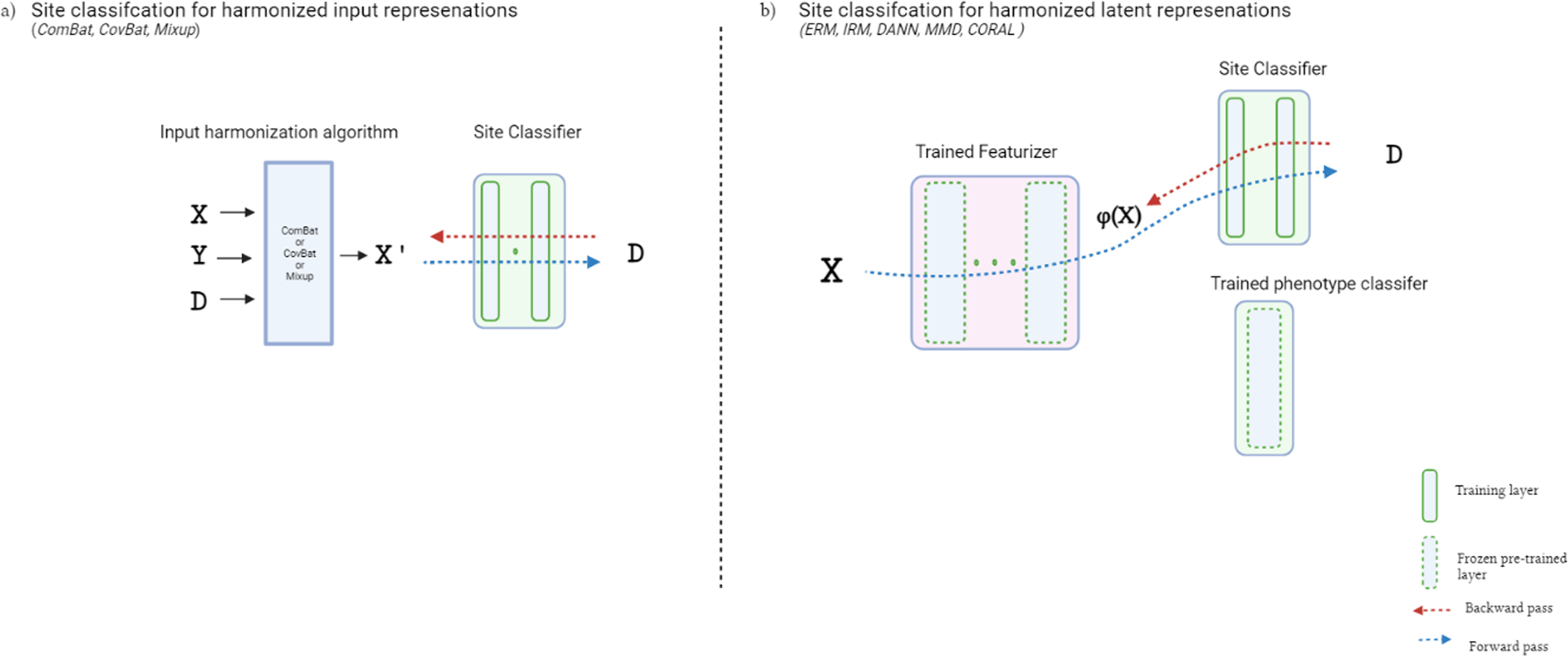
Pipeline to evaluate site classifcation accuracy of harmonized/baseline representations.

To answer the second question, we used two evaluation scenarios for the label of interest. The first is when there is access to data from all sites during training and the objective is to utilize the labelled data to build a model that generalizes on unseen data from the same sites. This is evaluated using a site-stratified k-fold validation scheme and we refer to it as the *domain adaption* problem. The second scenario is when we only have access to some sites during training and the objective is to train a model to generalize to unseen sites. This evaluated using a leave-one-site-out (LOSO) validation scheme and we refer to it as the *domain generalization* problem. Domain generalization is typically more challenging as there is no access to the test-set data distribution during training, yet, it represents a more realistic clinical scenario when the trained models are to be deployed on new unseen sites.

To ensure a fair comparison for all the methods, we followed a rigorous framework that accounts for variability that might arise due to hyperparameter configuration, model selection (i.e. when to stop training and which model to select for the prediction on the test set), and training stochasticity (randomness due to models initialization and data splits). To find the optimal hyperparameters for each method, we conducted a random search of 20 trials over the hyperparameter distribution of each method. We fixed the hyperparameter search distributions between the methods if possible barring some methodspecific hyperparameters. The best hyperparameter configuration (measured as the configuration yielding the highest test scores on a independent validation) were then used to train the final model and report the test results. For model selection during training and to stop training, we used an inner-validation split of (85% train - 15% validation) to monitor the convergence of the training and select the model for testing.

In reporting the test metrics for each method, we ran the entire experiment three times using three different seeds, thus making every random choice anew: hyperparameters random sampler, weights initializations, and dataset innerand outer-splits. Every number we report is a mean over these repetitions together with their estimated standard error.

## 4. Results

### 4.1. Post-harmonization site classification

We present site classifcation results after harmonization as described in Section 3. Figure 4 shows the test accuracy of the site classifcation experiment on the harmonized inputs/features on the largest four sites of the Rest-Metamdd dataset using static FC representation. The 5-fold accuracy dropped from 64.8*±*3.9% pre-harmonization to approximately chance level after data harmo-nization using the adopted methods described in Section 2.4. These empirical results suggest that the harmonization methods successfully removed site-effects that were originally found in the imaging derivatives and that drove high classification accuracies when predicting acquisition site. We observed similar behaviour for the ABIDE dataset and the dynamic representations of the data (See supplementary materials). Most interestingly in the results is that that latent features learned by the baseline model with no explicit harmonization contained no predictive information of the acquisition site, resulting in chance level performance in the site classification experiment akin to methods that explicitly harmonize the data/features. We discuss in Section 5 why this behaviour might occur.

**Figure 4:**
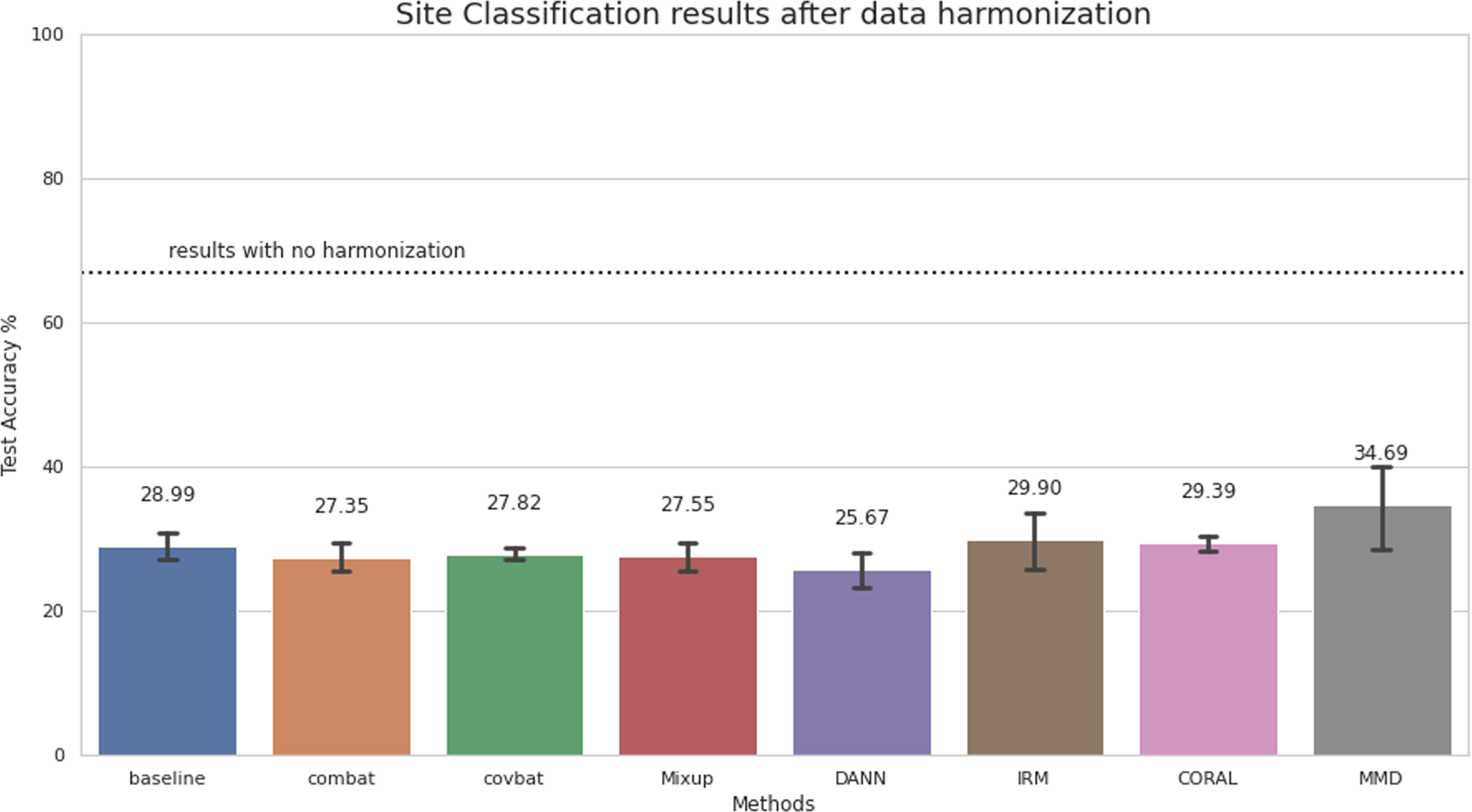
Site classification results after harmonizing the data for Rest-Meta-MDD using static representations. Chance level is 25%.

### 4.2. Domain adaptation

In this section, we present and compare the test results for the DL models optimized for the diagnosis of psychiatric disorders using a baseline model and harmonization methods in the domain adaptation scenario. To reiterate, domain adaption refers to the setup where there is availability of a number of samples from all the sites during training time. At inference, no data from unseen sites are presented to the model. We used site-stratified k-fold validation for reporting domain adaption results to ensure that all the data points are considered for testing and that all the sites are present at training and test time. The results in Figure 5 show the test performance for both the Rest-Meta-MDD and ABIDE datasets using static and dynamic representations when trained using different explicit harmonization techniques versus a baseline model trained on the unharmonized inputs. The results show that for all representation-dataset configurations, no harmonization method outperformed the baseline more than one accuracy point. Note that our results are generally on-par with the current reported state-of-the art performances for both datasets [37, 31, 38, 25, 39, 40, 41, 42].

**Figure 5:**
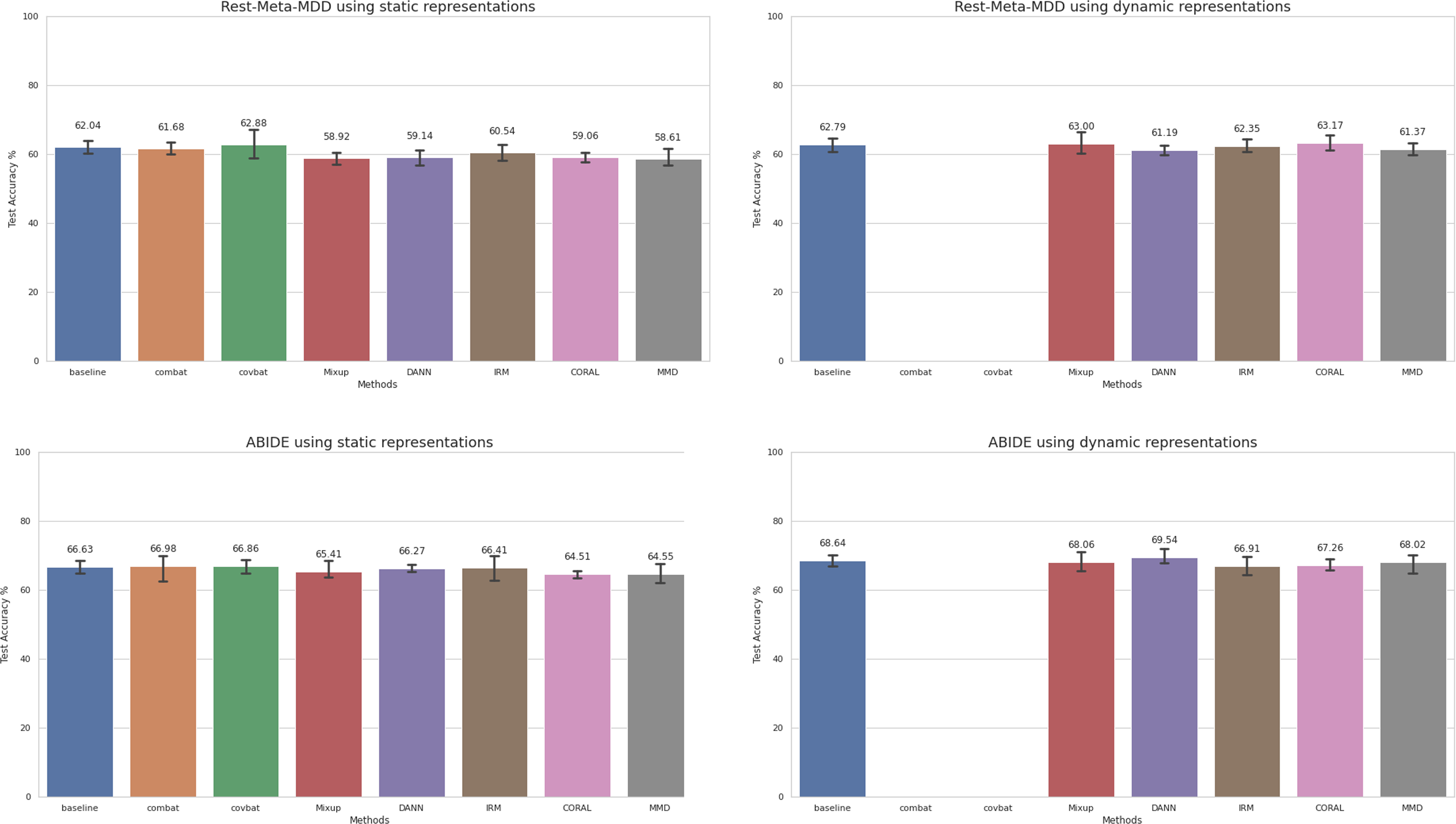
Domain adaptation results for different harmonization methods and the baseline on static and dynamic representations of Rest-Meta-MDD and ABIDE datasets. Chance level is 50%.

### 4.3. Domain generalization

We present the test results for the DL models optimized for the diagnosis of psychiatric disorders using a baseline model and harmonization methods in the domain generalization scenario. Tables 2-5 show the test accuracy for each leftout-site in both datasets using both static and dynamic representations. Again, the average LOSO accuracy for all dataset/representation combination showed no superior metric for any of the harmonization methods over the baseline. Interestingly, the average LOSO accuracy is on-par with the k-fold results even though in the LOSO setup, the model has no access to data from the test site during training. This behaviour further confirms that the models learn site-invariant features and the performance does not drop due to introduced heterogeneity from unseen sites.

**Table 2:**
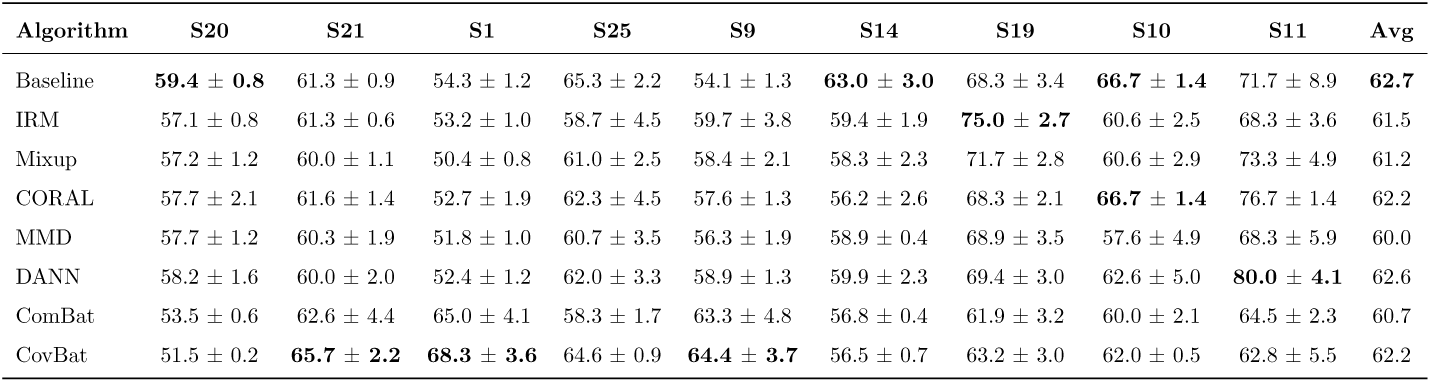
Rest-meta-MDD domain generalization results using static representations.

**Table 3:**
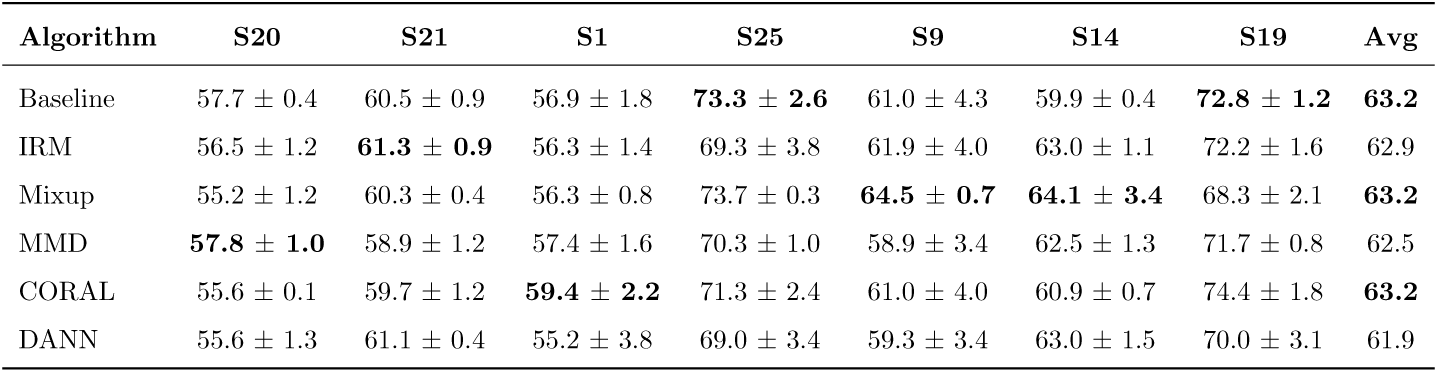
Rest-meta-MDD domain generalization results using dynamic representations.

**Table 4:**
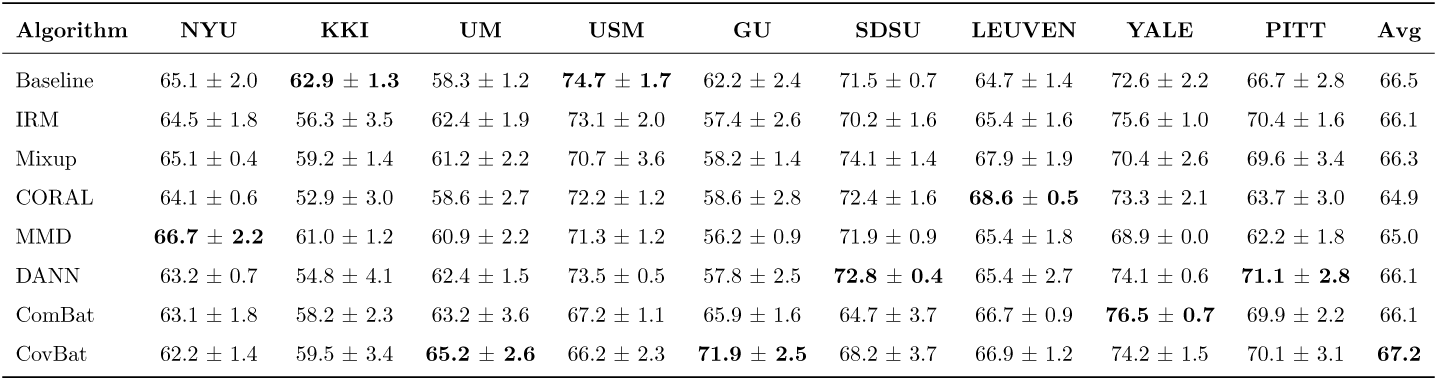
ABIDE domain generalization results using static representations.

**Table 5:**
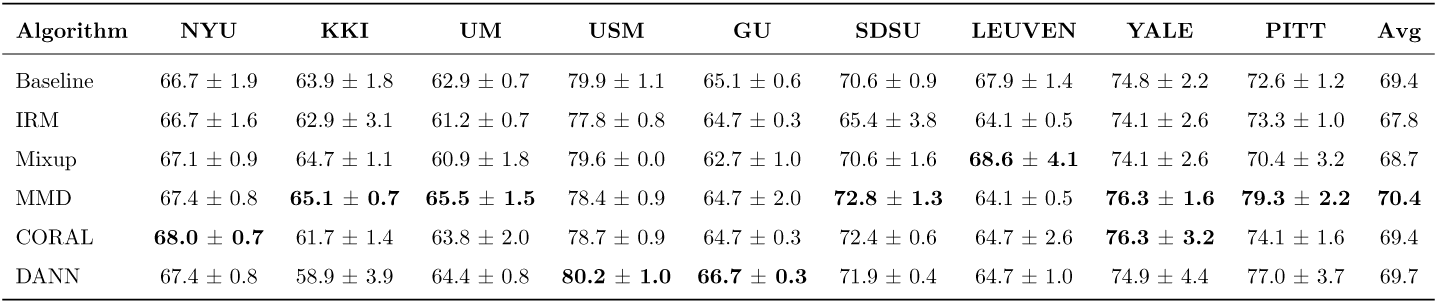
ABIDE domain generalization results using dynamic representations.

### 4.4. Further exploration

While the empirical results show that the studied harmonization methods did not improve the final outcome in domain generalization and domain adaption settings under our experimental setup, they open the door for more questions about other factors that can influence the results. Here we zoom into three factors that often vary across multi-site ML studies and can potentially lead to different conclusions about the efficacy of harmonization methods. Namely we study the implications of learning algorithm selection, number of sites within the dataset, and intra-site class imbalance. We conducted experiments for each variable with and without harmonization. For the sake of concision, we selected the ComBat technique in the experiments as a proxy for all the harmonization methods as it is currently the most popular approach within the field to harmonize the data.

#### 4.4.1. Effect of ML model choice & number of sites on generalization

In our experiments, we have used DL models to evaluate the efficacy of harmonization techniques on performance of diagnosis prediction. A natural question then arises, does the choice of learning model influence the effect of data harmonization? A possible explanation to the performance of the baseline is that DL models encode the data into a lower-dimensional latent feature space where site-discriminative features are lost. In this case, would a more simple linear model such as a logistic regression classifier benefit from harmonizing the data?

Another variable of interest is the number of acquisition sites in the dataset. Intuitively, as the number of sites in the dataset grow, the risk that a learning model optimized to minimize the disorder prediction error would over-fit on site information diminishes. In the experiments conducted thus far, we used nine sites (each with N *≥* 40) to evaluate the generalizability of the performance. This is however typically not the case for the majority of multi-site datasets where the number of sites is much smaller. Our setup could thus under-estimate the efficacy of harmonization in such datasets.

To account for these potential confounding variables in our framework, we compared the performance of a logistic regression classifier with and without combat harmonization on the tasks of MDD and ASD diagnosis from static FC in the domain generalization setup. Further, we conducted the experiments using different numbers of sites *M* ∈ [3,11], where for each *M* the experiments are repeated 10 times each using randomly sampled sites. We report the results in Figure 6. Our results show that the LR model marginally benefited from combat harmonization at all sampled *M* configurations on the ABIDE dataset while the pattern is not clear for MDD-rest where harmonization appears to benefit the model at low *M* and then the baseline eventually catches up as *M* increases beyond five sites. A notable observation from this experiment orthogonal to harmonization is that test accuracy does not improve much as the number of sites in the training dataset increases (and hence the number of training samples). We suspect this could be due to the noise in the data and clinical labels which create an upper bound on the performance of the learning model at the current order of magnitude of available training samples.

**Figure 6:**
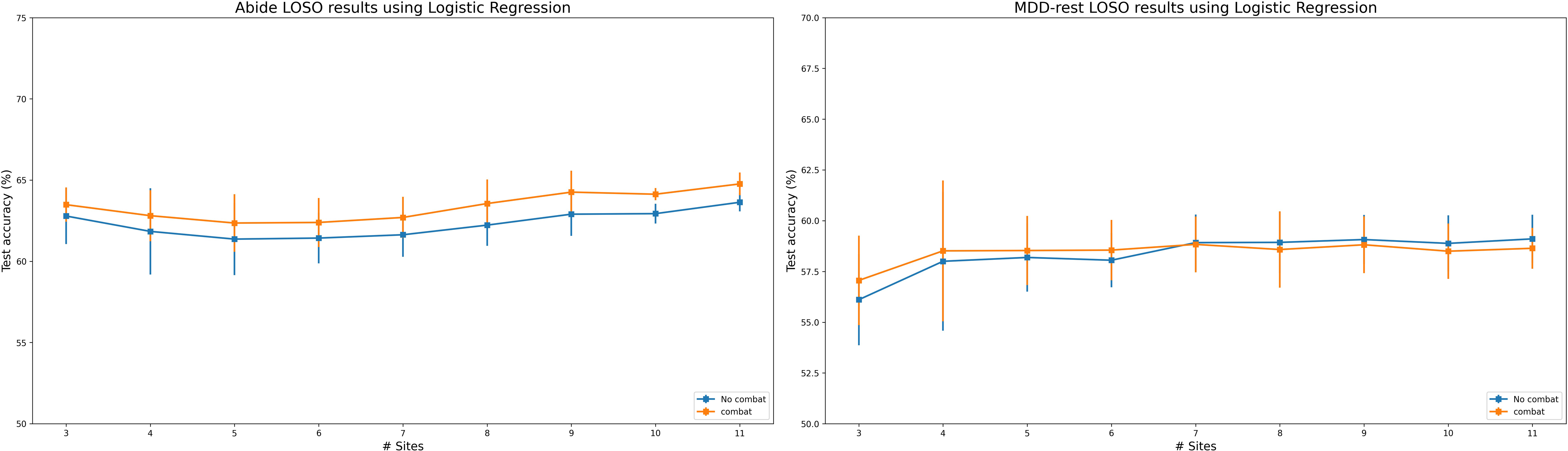
Domain generalization results with and without ComBat harmonization using logistic regression and variable number of sites during training on both datasets on static FC representation. Chance level is 50%.

### 4.5. Effect of intra-site class imbalance on generalization

A topic of controversy in neuroimaging multi-site studies, is that disease classifiers can overfit on site information if the class distribution across sites is highly non-uniform, resulting in unrealistic evaluation of the model performance. This is even more pronounced when the label of interest is challenging for the model and it is easier to predict the site to minimize its prediction error. This is often the case with psychiatric ML studies if researchers fail to account for high intra-site class imbalance either in the objective function of the model or data sampling. The datasets used in this study have a nearly uniform class distribution across most of the sites (see Figure 1) and in our experiments, we ensured a perfect uniform class distribution by under-sampling the majority class within each site during training. In this section, we aimed to investigate the effect of harmonization on model performance under different intra-site class distributions. Ideally, for datasets with highly class-imbalanced sites, harmonization methods should prevent erroneous convergence, represent a fair evaluation of the model, and even enable the use of the full dataset without under-sampling. To investigate this, we designed an experiment where class distribution is controlled by a parameter *α* ∈ [0.5, 1]. When sampling data from each site, we sampled class A with a probability *α*, and class B is thus sampled with probability 1 *− α*. Given that the base distribution in our sites is nearly uniform, we can thus control the class distribution by *α*. The class balance of the entire dataset is maintained by alternating classes A and B at each site. (i.e, under sampling class A at site 1, under sampling class B at site 2, and so on.). We present the results for both MDD and ASD classifcation on the MLP at different *α* in Figure 7. The results confirm that for unharmonized data, increasing intra-site class imbalance linearly increases the performance of the model from the baseline results observed throughout all our experiments. This indicates that at higher *alpha* the model converges to learning site differences instead of the biological label of interest. But when using ComBat, the performance does not diverge with class imbalance, indicating the learning is not confounded at high intra-site class imbalance.

**Figure 7:**
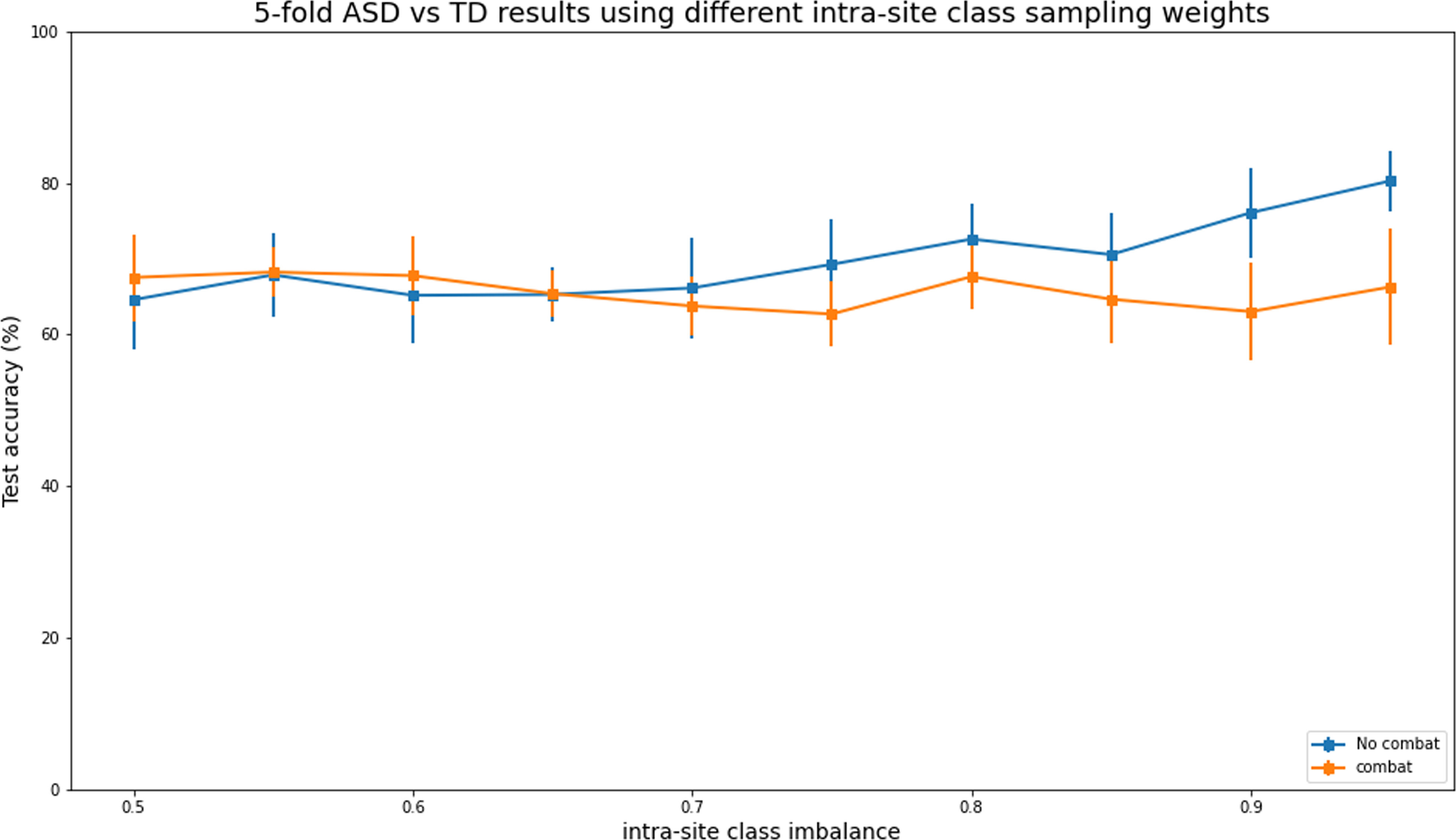
Domain adaptation ABIDE results (5-fold balanced accuracy) with and without ComBat harmonization under different intra-site class distributions. Increasing intra-site class imbalance causes the model with unharmonized inputs to diverge from learning diseasediscriminative features to site-discriminative features. Applying ComBat harmonization on the input prevents this behaviour and stabilize learning under high intra-site class imbalance setting.

## 5. Discussion

The goal of this work was to evaluate the efficacy of applying harmonization techniques on the performance of learning models to diagnose psychiatric disorders from rs-fMRI data. Specifically, we compared the performance of seven different popular harmonization techniques against an equally powerful baseline on static and dynamic representations of fMRI data for the tasks of diagnosing MDD vs HC and ASD vs TD in two publicly available multi-site datasets using deep learning models. The main finding of this work is that while harmonization techniques successfully remove site-effects in the data, they do not improve the final outcome on the desired tasks against the baseline for both static and dynamic representations of the data and in both domain adaptation and domain generalization settings. Investigating the behaviour of the baseline showed that when then model is optimized solely for the task of disease diagnosis and intrasite class distribution is balanced, it learns site-invariant feature representations which could well explain the empirical performance against explicit harmonization methods. Further investigative experiments showed no effect of the number of sites in the dataset or chosen learning algorithm on this conclusion. A key insight however is the advantage of harmonization in experimental setups with high inter-site class imbalance, where harmonization methods reduce the risk of the disease classifier diverging from learning the desired task. We observed this behaviour with unharmonized data: where intra-site class imbalance increased, the tendency of the model to learn spurious site features, and hence reporting erroneous inflated results increased. Harmonization methods could thus be used to “combat” this behaviour and ensure that the reported results are that of the biological label of interest. For the remainder of the discussion, we examine the empirical results from a theoretical perspective, i.e. why does the baseline model learn site-invariant features? Further we highlight some of the common pitfalls in harmonization studies. Finally we make some remarks about our framework including its utility to the community and some of the limitations of our problem setup.

### Why does the baseline model learn site-invariant features?

The need for harmonization stems from the fact there exists a distributional shift within the training dataset or between the training and test sets. As we have seen in our site experiment, data distribution *P* (*X*) is indeed different between sites and one can easily train a classifier to identify the source of the distribution. A special case however, is when the joint distribution between domains is different *P_A_*(*Y, X*) ≠ *P_B_*(*Y, X*) but the conditional distribution is similar *P_A_*(*Y |X*) = *P_B_*(*Y |X*). This is commonly referred to in the literature as *covariate shift* and also encapsulates transformations to the data *P_A_*(*Y |ϕ*(*X*)) = *P_B_*(*Y |ϕ*(*X*)[43]. The goal of training supervised ML classifiers is to find such transformations of the data *ϕ*(*X*) that maximizes the log likelihood of probability of the correct label. If there is indeed a conditional independence between the label and domain, an optimal model trained to minimize the empirical risk of prediction error of the label has no incentive to learn a transformed feature space which contains spurious site information [44].

In one often cited example, [45] trained a convolutional neural network to classify camels from cows. In the training dataset, most of the pictures of cows had green pastures, while most pictures of camels were in the desert. The model picked up the spurious correlation and associated green pastures with cows thus failing to classify pictures of cows on beaches correctly. This is an example of covariate shift, as the casual features of the label we are interested should be independent of where the picture is taken. However because the training dataset is biased, the model fails to learn the desired causal features. This is simply fixed by collecting a dataset that contains diverse examples or by data augmentation. This behaviour has been observed in multiple domains [46, 47, 48, 22] where advanced harmonization methods did not improve on baseline methods when a suitable learning algorithm is chosen and the datasets are unbiased towards the domain.

We believe that this is similar to the case in our setup. Despite strong site-effects, because the datasets are balanced and the label is orthogonal to the site, the results suggest that the baseline model learns site-invariant features as seen in Section 4.1 and thus eliminating the need for explicit harmonization methods. This is not the case however in unbalanced sites where the baseline exhibit similar behaviour as in [45] and learns spurious domain features.

### Common pitfalls in evaluating harmonization methods

As we alluded earlier, data harmonization is a major area of controversy in the neuroimaging community. In this section we review some of the common pitfalls that hinders the reproducability of harmonization methods across research groups and fuels this controversy.

#### Under-optimized baselines

A number of harmonization studies typically spend time and computational resources on optimizing a proposed harmonization algorithm, and do not allocate the same resources to build an equally powerful baseline model. For example, in our experiments, we observed that evaluating a baseline before training convergence or without the use of an inner validation to avoid overfitting, hampers its generalization performance on the test set. Other examples include training the baselines off-the-shelf without any search for optimal hyperparameters, while doing a exhaustive hyperparameter search for the models used in harmonization experiments. The role of ML practitioners is to ensure a fair comparison for all the methods for a faithful evaluation of the efficacy of harmonization methods.

#### Inconsistencies in experimental conditions & model selection

Similarly, experimental conditions should not vary between the methods. [22] finds that model selection (when to stop training) plays an important role in evaluating domain generalization algorithms and recommends explicit specification and standardization of model selection method for any harmonization algorithm. Further, the experiments should be run using several random seeds and data splits should be consistent between the methods.

#### Data Leakage

Several open-source online implementations of harmonization methods utilize the entire dataset including the test samples to harmonize the data. This behaviour is a concrete example of double dipping. This does not only leak test-data statistics into training, but for some harmonization methods where the label of interest is required, it also leaks the test labels. This often leads to unrealistic optimistic results only due to the leakage and not because of data harmonization.

### The utilities and limitations of our framework

To conduct the experiments reported in this work, we have built an open-source framework on top of the Domain-bed[22] library to ensure standardization of architectures/experimental setups and enable the reproducability of the results. While we have studied harmonization on two datasets with two different clinical targets, our framework can easily be extended to any fMRI dataset and for any phenotype using a few lines of code. This also includes integrating different representations of the data (e.g. 4D volumes, Graphs) along with any custom neural network architectures. We encourage researchers working on this topic to include their harmonization methods or architectures in our framework for evaluating the efficacy of new proposed methods against existing solutions.

While the empirical results in our work guide suggest that harmonization algorithms do not improve the generalization performance of fMRI phenotype prediction models, it remains possible that harmonization may be useful in other settings than those we have investigated. And although our results suggest that deep learning models appear to handle site heterogeneity intrinsically by learning site-invariant features, a small benefit of harmonization was observed for logistic regression for one of the two datasets. The main use for harmonization appears to be the situation with class-imbalance across sites, when algorithms could otherwise use site information to predict the label of interest. And although harmonization did not improve disorder prediction, it did not reduce performance either. Correct fMRI data harmonization therefore does not appear to come at a costs. However, we identified several situations where incorrect implementation of data harmonization could result in erroneous conclusions. We therefore advice to use fMRI data harmonization primarily to combat class imbalance, rather than incorporating it as standard strategy for multi-site studies.

